# Cell-free styrene biosynthesis at high titers

**DOI:** 10.1101/2020.03.05.979302

**Authors:** William S. Grubbe, Blake J. Rasor, Antje Krüger, Michael C. Jewett, Ashty S. Karim

## Abstract

Styrene is an important petroleum-derived molecule that is polymerized to make versatile plastics, including disposable silverware and foamed packaging materials. Finding more sustainable methods, such as biosynthesis, for producing styrene is essential due to the increasing severity of climate change as well as the limited supply of fossil fuels. Recent metabolic engineering efforts have enabled the biological production of styrene in *Escherichia coli*, but styrene toxicity and volatility limit biosynthesis in cells. To address these limitations, we have developed a cell-free styrene biosynthesis platform. The cell-free system provides an open reaction environment without cell viability constraints, which allows exquisite control over reaction conditions and greater carbon flux toward product formation rather than cell growth. The two biosynthetic enzymes required for styrene production were generated via cell-free protein synthesis and mixed in defined ratios with supplemented L-phenylalanine and buffer. By altering the time, temperature, pH, and enzyme concentrations in the reaction, this approach increased the cell-free titer of styrene from 5.36 ± 0.63 mM to 40.33 ± 1.03 mM, an order of magnitude greater than cellular synthesis methods. Cell-free systems offer a complimentary approach to cellular synthesis of small molecules, which can provide particular benefits for producing toxic molecules.

**Highlights:** A cell-free system for styrene biosynthesis was established. This *in vitro* system achieved styrene titers an order of magnitude greater than the highest reported concentration *in vivo.*

## 1. Introduction

Metabolic engineering has enabled the production of commodity chemicals and valuable small molecules by genetically modifying microorganisms and overexpressing heterologous enzymes (Keasling, 2010; Liu and Nielsen, 2019; Nielsen, 2001; Stephanopoulos, 1994; Tyo et al., 2007). Target biochemicals, such as butanol (Shen and Liao, 2008) and mevalonate (Martin et al., 2003), are often selected based on their utility for society. An important large-volume commodity chemical is styrene, which is produced globally on the scale of 30 million tons per year; 60% of the product is utilized for molded or foamed polystyrene and the remainder contributes to industrially important copolymers, such as styrene-acrylonitrile and styrene-butadiene (James and Castor, 2011). However, styrene production is an entirely petroleum-derived process that requires large excesses of steam and is responsible for over 100 million tons of greenhouse gas emissions each year (Wu et al., 1981; Zheng and Suh, 2019). Through metabolic engineering of *Escherichia coli*, biosynthesis of styrene from glucose via the shikimate pathway was demonstrated as a potential, sustainable alternative to traditional styrene synthesis, albeit at low titers up to ~2.6 mM styrene (McKenna and Nielsen, 2011). Recent efforts utilized genome editing and solvent extraction techniques to increase styrene titers in *E. coli* cultures, but the maximum concentration only reached ~3.4 mM (Liu et al., 2018). Clearly, the cellular toxicity of styrene greatly limits biosynthesis titers and the feasibility of commercial styrene biosynthesis (Araya et al., 2000). Circumventing cellular toxicity could prove useful for the biochemical production of styrene.

Recent advances in cell-free technologies have showcased their utility for studying biological processes and engineering biological systems (Bogorad et al., 2013; Garenne and Noireaux, 2019; Jaroentomeechai et al., 2018; Kightlinger et al., 2019; Lee et al., 2019; Martin et al., 2018; Schwander et al., 2016; Silverman et al., 2019). For example, several studies have shown that crude extracts contain native metabolic enzymes and cofactor regeneration responsible for robust cell-free protein synthesis (Caschera and Noireaux, 2014; Des Soye et al., 2019; Jewett et al., 2008; Jewett and Swartz, 2004) and activation of key metabolic reactions in the cell-free environment (Dudley et al., 2015; Jewett and Swartz, 2004; Karim et al., 2018; O’Kane et al., 2019). In fact, cell-free systems have been used for biosynthesis of a wide variety of molecules, including 2,3-butanediol (Kay and Jewett, 2015), mevalonate (Dudley et al., 2016), natural products (Goering et al., 2017; Liu et al., 2019; Zhuang et al., 2020), *n*-butanol (Karim and Jewett, 2016; Krutsakorn et al., 2013; Reisse et al., 2016), terpenes (Dudley et al., 2019; Korman et al., 2017), and polyhydroxyalkanoates (Kelwick et al., 2018) using processes with purified enzymes or crude cell extracts (Claassens et al., 2019; Rollin et al., 2018). Cell-free systems provide an open reaction environment and rapid design-build-test cycles to reconstitute biosynthetic pathways *in vitro* to compliment and inform metabolic engineering efforts in cells (Bundy et al., 2018; Dudley et al., 2015; Gregorio et al., 2019; Hodgman and Jewett, 2012; Karim et al., 2019a). More importantly, cell-free systems have shown improved tolerance to toxic small molecules compared to living systems (Kay and Jewett, 2019), providing evidence that cell-free biomanufacturing platforms may be advantageous when cellular systems prove impractical.

In this work, we established a cell-free platform for styrene biosynthesis to increase the achievable titer in biological systems by circumventing the toxicity limits of styrene *in vivo*. We constructed this system in two parts. First, we used cell-free protein synthesis in *E. coli* crude extracts to express the two non-native enzymes required to convert L-phenylalanine (L-Phe) to styrene: phenylalanine ammonia lyase 2 (PAL2) from *Arabidopsis thaliana* and ferulic acid decarboxylase 1 (FDC1) from *Saccharomyces cerevisiae* (Liu et al., 2018; McKenna and Nielsen, 2011). Next, we combined these extracts enriched with biosynthetic enzymes with L-Phe and buffer to produce styrene *in vitro* (**Figure 1**). We further optimized the cell-free system by tuning enzyme ratios, reaction temperature, and reaction pH of these reactions to reach styrene titers over 40 mM, an order of magnitude greater than previous cellular efforts. We anticipate this work will expand the application space of cell-free systems and spur new research efforts in the metabolic engineering of toxic chemicals.

**Figure 1.**
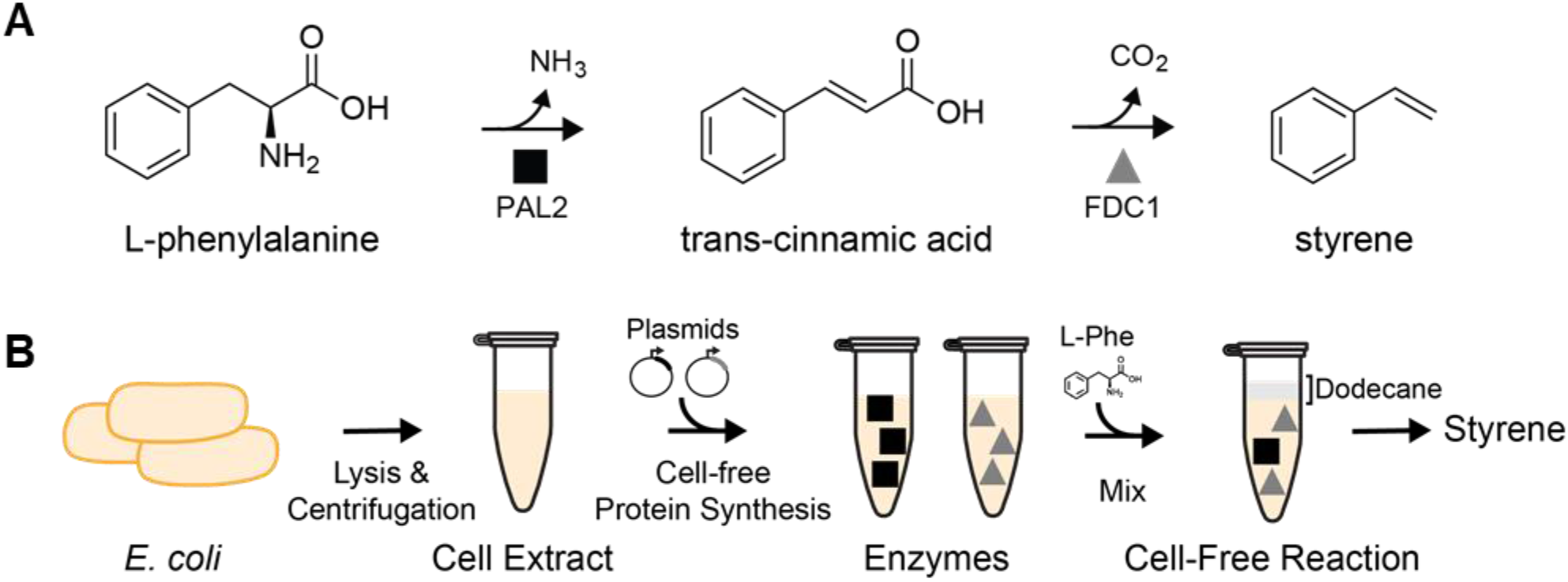
Enzymatic conversion of L-phenylalanine to styrene. **(A)** A two-step conversion is catalyzed by phenylalanine ammonia lyase (PAL2) and ferulic acid decarboxylase (FDC1) to produce styrene with ammonia and carbon dioxide as byproducts. **(B)** The enzymes were synthesized with *E. coli* cell extract and mixed with phenylalanine in quantifiable proportions.

## 2. Materials and Methods

### 2.1. Bacterial strains and plasmids

Plasmid propagation was performed in *E. coli* DH5α (NEB), and *in vitro* enzyme expression was performed in cell extract prepared from *E. coli* BL21 Star (DE3) (Life Technologies). Plasmid pSpal2At, containing the PAL2 gene from *Arabadopsis thanliana*, was a gift from David Nielsen (Addgene plasmid 78286). This gene was cloned into the pJL1 plasmid (Addgene plasmid 69496) using Gibson Assembly after PCR amplification with oligonucleotides from IDT (5’-tttaagaaggagatatacatATGGATCAAATCGAAGCAATG-3’ and 5’-tttgttagcagccggtcgacTTAGCAAATCGGAATCGG-3’). The FDC1 gene from *Saccharomyces cerevisiae* was synthesized and cloned into the pJL1 plasmid for expression by Twist Biosciences. Sequences are provided in **Supplementary Table 1**. Propagated plasmids were purified using the ZymoPURE Plasmid Miniprep Kit (Zymo Research).

### 2.2. Cell extract preparation

*E. coli* extracts were prepared as previously described.(Karim and Jewett, 2018; Kwon and Jewett, 2015) In brief, BL21 Star (DE3) cells (Life Technologies) grown in 1 L of 2xYTPG media in full-baffle shake flasks at 37°C. At an OD_600_ of 0.4, 1 mM of IPTG was added to induce T7 RNA polymerase production. Cells were harvested at an OD_600_ of 3.0. Cells were pelleted via centrifugation at 5,000g for 10 minutes at 4°C, washed three times with cold S30 buffer (10 mM tris acetate, pH 8.2; 14 mM magnesium acetate; 60 mM potassium acetate; and 1 mM dithiothreitol), flash-frozen with liquid nitrogen, and stored at −80°C. For lysis, cells were thawed on ice and resuspended in 1 mL of S30 buffer per gram wet cell mass and then lysed in an EmulsiFlex-B15 homogenizer (Avestin) in a single pass at a pressure of 20,000-25,000 psi. Cellular debris was removed by two rounds of centrifugation at 12,000g for 30 minutes at 4°C, and the final supernatant was flash-frozen with liquid nitrogen and stored at −80°C until use.

### 2.3. Cell-free protein synthesis (CFPS) reactions

CFPS reactions for *in vitro* production of enzymes were assembled with 6 nM template DNA, 10 mg/mL *E. coli* extract, and the cofactors and crowding agents in 57 mM HEPES buffer. These reactions contained 8 mM magnesium glutamate; 10 mM ammonium glutamate; 130 mM potassium glutamate; 1.2 mM adenosine triphosphate; 0.85 mM each of guanosine, uridine, and cytidine triphosphates; 0.034 mg/mL folinic acid; 0.171 mg/mL transfer RNAs; 33.33 mM phosphoenolpyruvate; 2 mM of all 20 canonical amino acids; 0.40 mM nicotinamide adenine dinucleotide; 0.27 mM cofactor A; 1 mM putrescine; 1.5 mM spermidine.(Jewett and Swartz, 2004) The expression level of each enzyme was quantified using radioactive leucine incorporation assays as previously described.(Jewett et al., 2008) All reagents and chemicals were purchased from Sigma-Aldrich unless otherwise specified.

### 2.4. Cell-free metabolic engineering (CFME) reactions

Styrene biosynthesis reactions contained 8 mM magnesium glutamate, 10 mM ammonium glutamate, 134 mM potassium glutamate, 100 mM BisTris buffer, 0.5 mM kanamycin, varying concentrations of PAL2 and FDC1 enzymes from CFPS ranging from 0.05 to 1 μM, and 25 or 50 mM L-Phe. A layer of dodecane was placed atop the reaction to capture volatile styrene.(Dudley et al., 2019)

### 2.5. Metabolite analysis

Styrene was quantified by diluting 20 μL of dodecane overlay into 200 μL of ethyl acetate containing 0.5 mM *trans*-caryophyllene (Sigma) as an internal standard. 1 μL of this mixture was injected into an Agilent 7890A Gas Chromatograph with 5977A MSD (Agilent, Santa Clara, CA) using an Agilent HP-5MS (30 m length x 0.25 mm i.d. x 0.25 μm film) column with helium carrier gas at constant flow of 1 mL·min-1. The inlet temperature was 70 °C and initial column temperature held at 70 °C for 1 minute, increased at 25 °C·min-1 to 250 °C, and maintained at 250 °C for 3 minutes. The injection volume was 1 μL with a split ratio of 20:1. Extracted ion chromatograms (EIC) for 104 m/z (styrene, peak at 2.97 min) and 133 m/z (caryophyllene, peak at 6.34 min) were integrated using Agilent MassHunter Quantitation Analysis software. Concentrations were determined by use of a standard curve (**Supplementary Figure S1**) generated by comparison to styrene (Sigma) standards mixed in dodecane with mock cell-free reactions containing green fluorescent protein in place of the biosynthetic enzymes that were incubated for 24 hours (Dudley et al., 2019).

### 2.6. pH measurements

Samples were analyzed with a Thermo Scientific™ Orion™ ROSS Ultra™ Refillable pH/ATC Triode™. Reactions for which a pH was set prior to reaction start were measured with a mixture of all components except the enzyme-enriched CFPS reactions to avoid premature reaction initiation. Reaction pH was adjusted with glacial acetic acid or 5 N KOH as necessary. Measurements of pH over time were taken after sampling reactions for metabolite analysis.

## 3. Results & Discussion

To establish a cell-free platform for styrene biosynthesis, we took a two-pronged approach: first establishing enzyme synthesis and pathway assembly, and second optimizing physiochemical conditions for improved production.

### 3.1. Enzyme synthesis and styrene pathway assembly

We first demonstrated the ability to express functional enzymes for the styrene biosynthesis pathway *in vitro* and to reliably capture the volatile styrene product. Cell-free protein synthesis (CFPS) enables rapid production of the enzymes for styrene biosynthesis, PAL2 and FDC1 (**Figure 2A-B**). Using CFPS, we produced 4.99 ± 0.36 μM soluble PAL2 and 5.93 ± 0.47 μM soluble FDC1 over the course of a 20-hour reaction at 30 °C. In an attempt to express greater soluble fractions of PAL2 and FDC1, we decreased the temperature of the reactions from 30 °C to 16 °C. The decreased temperature increased enzyme solubility at 6 h from ~70% to ~80% for PAL2 and from ~60% to ~67% for FDC1. For both temperatures, a majority (~80%) of the soluble protein made during the reaction is produced by 6 h. Therefore, we chose to run CFPS reactions at 16 °C and stop reactions at 6 h to accelerate the workflow while still obtaining sufficient concentrations of soluble enzymes for all subsequent reactions.

**Figure 2.**
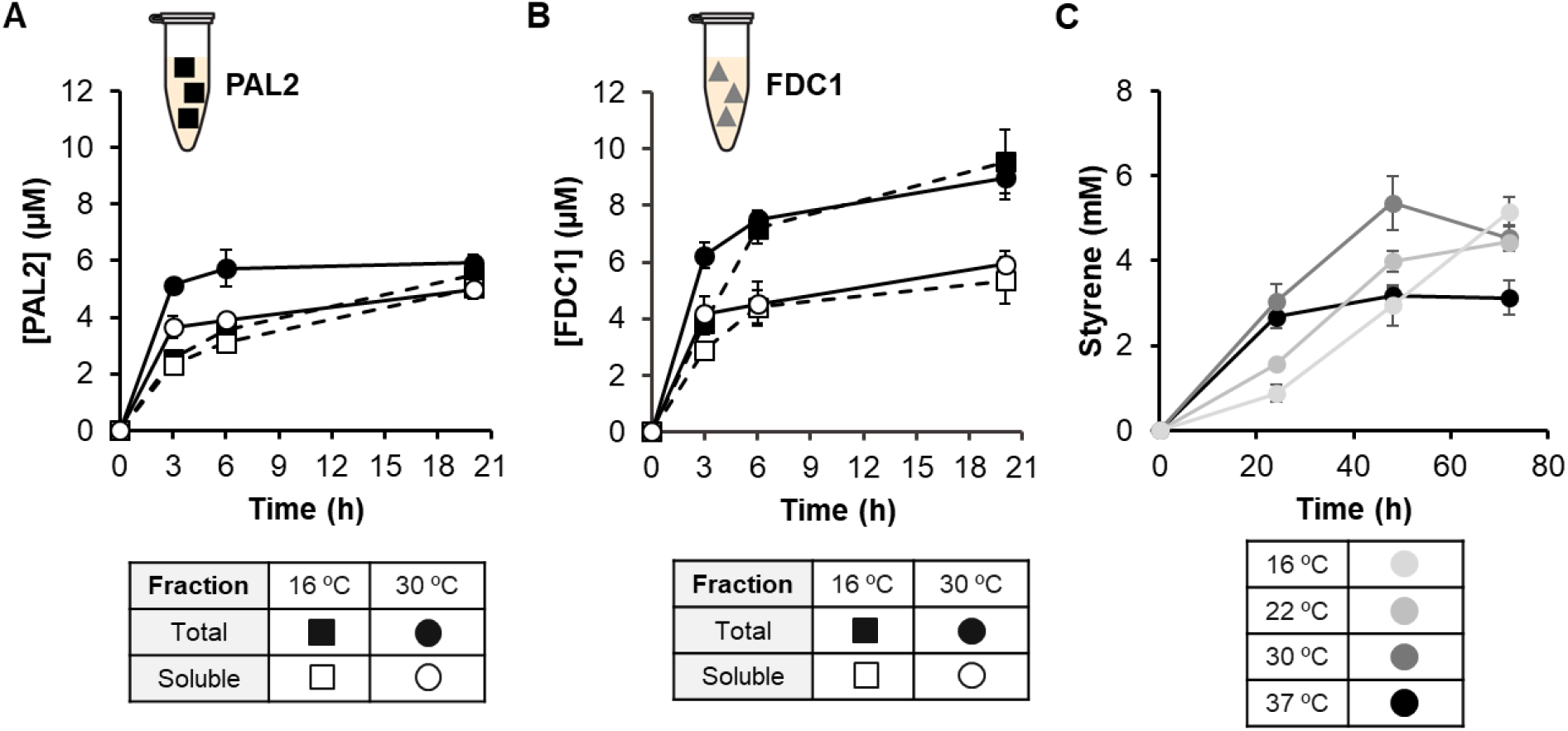
Expression and activity of the styrene biosynthetic pathway. Expression of PAL2 **(A)** and FDC1 **(B)** was assessed by radioactive leucine incorporation, and soluble enzyme fractions were used for subsequent quantification of enzyme concentrations. **(C)** Combining 0.5 μM PAL2 and FDC1 enabled styrene production, and the maximum titer of 5.36 mM was produced after 48 hours at 30 °C. Error bars represent standard deviation (A-B) or standard error (C) of 3 technical replicates.

To test for activity, we ran several reactions measuring phenylalanine conversion and styrene production. We confirmed that cell-free expressed PAL2 alone converts L-Phe to trans-cinnamic acid (**Supplementary Figure S2A**) and PAL2 combined with FDC1 produces styrene (**Supplementary Figure S2B**) as determined by HPLC. Additionally, no styrene is produced in cell-free reactions lacking the exogenous enzymes and supplemental L-Phe (**Supplementary Figure S3**). Accurate quantification of styrene can be difficult due to its volatility and lack of solubility in aqueous media; therefore, we ran our reactions with a dodecane overlay to capture styrene and detect it by GC-MS. This approach is often used to extract volatile compounds from both *in vivo* (Liu et al., 2018) and *in vitro* (Dudley et al., 2019) systems. We found that larger ratios of overlay to reaction volume enabled greater styrene recovery relative to the *trans*-caryophyllene internal standard without inhibiting biosynthesis (**Supplementary Figure S4**).

After demonstrating enzyme activity and the ability to measure styrene, we investigated the best temperature during the biosynthesis segment of the reaction for producing styrene. To do so, we mixed PAL2 and FDC1 after cell-free expression in a second pot reaction at a final concentration of 0.5 μM each. We then incubated reactions containing 25 mM L-Phe at 16, 22, 30, and 37 °C (**Figure 2C**). The rate of styrene production was highest at 30 °C and produced a maximum titer of 5.36 mM styrene after 48 hours. This titer demonstrates a ~1.5-fold increase over the observed inhibitory concentration of styrene for *E. coli* (Liu et al., 2018), confirming the potential for this cell-free platform to produce toxic compounds.

### 3.2. Optimization of enzyme ratio and physiochemical conditions

We subsequently stepped through a series of optimizations to improve cell-free styrene synthesis by exploiting the open reaction environment which enables precise control over enzyme concentrations and physiochemical conditions (Karim et al., 2018; Karim and Jewett, 2016; Silverman et al., 2019). Consistent with our previous work, we first normalized the total volume of CFPS added to a biosynthesis reaction by supplementing with CFPS mixtures without plasmid. By doing this, we minimize detrimental effects that additional CFPS volume, specifically the additional small molecules, tends to cause (*e.g.*, decreased final titers of the desired product) (Karim et al., 2018). However, we also wanted test whether this is universally true or potentially specific to previously studied pathways. We found that biosynthesis reactions with increasing amounts of CFPS fraction result in increased styrene production (**Supplementary Figure S5**). Our reactions are less inhibited by the CFPS mixtures likely because we are observing a two-step biosynthesis from L-Phe rather than longer pathways that take advantage of glycolysis that are known to have competition with several pathways branching from pyruvate and acetyl-CoA. These results suggest that the impact of CFPS reagents should be investigated for each new biosynthetic pathway tested *in vitro*.

We next decided to tune the biosynthetic enzyme ratio by mixing different volumes of cell-free expressed PAL2 and FDC1 in the reactions. We ran 36 unique reaction conditions varying the final PAL2 and FDC1 concentrations from 0 to 1 μM (**Figure 3A**). The best condition produced up to 18.03 ± 2.34 mM styrene from 25 mM of added L-Phe. As hypothesized, styrene titer generally increased with increasing enzyme concentrations. However, the best 8 enzyme ratios all produced 16-18 mM styrene, which suggested substrate limitation may prevent higher titers. We doubled the initial concentration of added L-Phe to 50 mM and ran reactions using the top eight enzyme combinations in an attempt to further increase styrene yield. The best reaction condition produced 24.83 ± 0.66 mM styrene with 0.25 μM PAL2 and 1 μM FDC1 (**Figure 3B**). Increasing the substrate concentration enabled differentiation between the best conditions with 25 mM L-Phe in **Figure 3A**, but the modest 6-7 mM increase in product from 25 mM additional substrate indicated diminishing biosynthetic potential; thus, L-Phe concentrations greater than 50 mM were not examined.

**Figure 3.**
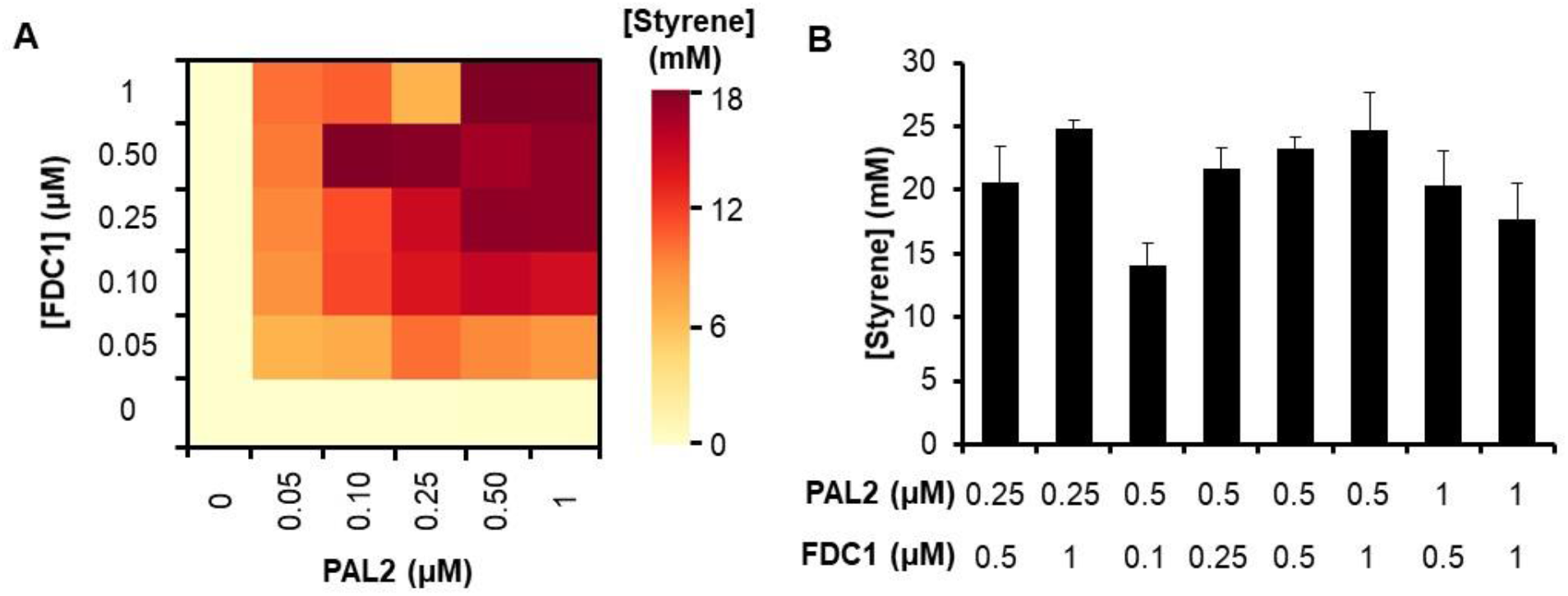
Modulating enzyme concentrations enhances styrene biosynthesis. **(A)** Increasing the concentration of each enzyme increased the final titer, as expected. **(B)** All conditions producing more than 16 mM styrene from 25 mM L-Phe were run with 50 mM L-Phe to identify the optimal enzyme ratio – 0.25 μM PAL2 with 1 μM FDC1. Error bars represent standard deviation of 3 technical replicates.

### 3.2. Optimization of physiochemical conditions

To further optimize the reaction environment for styrene biosynthesis, we explored a range of acidic and alkaline conditions due to the dramatic changes that pH can cause in cell-free systems (Calhoun and Swartz, 2005; Karim et al., 2019b). We tested six different initial pH conditions in reactions for styrene synthesis ranging from pH 5.6 to pH 9.5. Our initial condition began at pH 7.5-7.8 and ended near pH 8 (**Figure 4**; dark blue). We observed a strong pH dependence for styrene biosynthesis across the broad pH range, with a maximum titer of 40.33 ± 1.03 mM styrene after 72 hours between pH 7 and pH 8 (**Figure 4A**). The most alkaline reaction (pH 9.5) produced less than 1 mM styrene, whereas the most acidic reaction (pH 5.6) produced 3.27 ± 2.85 mM styrene – nearly equivalent to the highest reported titer from *in vivo* biosynthesis (Liu et al., 2018). Reaction rates remained steady over 3 full days, which was considerably slower than the reported catalytic rates of purified PAL2 and FDC1 (**Figure 4B**) (Cochrane et al., 2004; Payne et al., 2015). Although enzyme behavior can differ *in vitro* between purified systems and our crude cell extracts, the observed pH optimum for styrene biosynthesis lies within a reasonable range based on reported pH optima of 8.4-8.9 and 6.5 for PAL2 and FDC1, respectively (Cochrane et al., 2004; Lin et al., 2015). Additionally, the reaction pH changed very little over time without active glycolysis producing acetate and lactate, which provided a stable reaction environment without the need for a strong buffer (**Figure 4C**).

**Figure 4.**
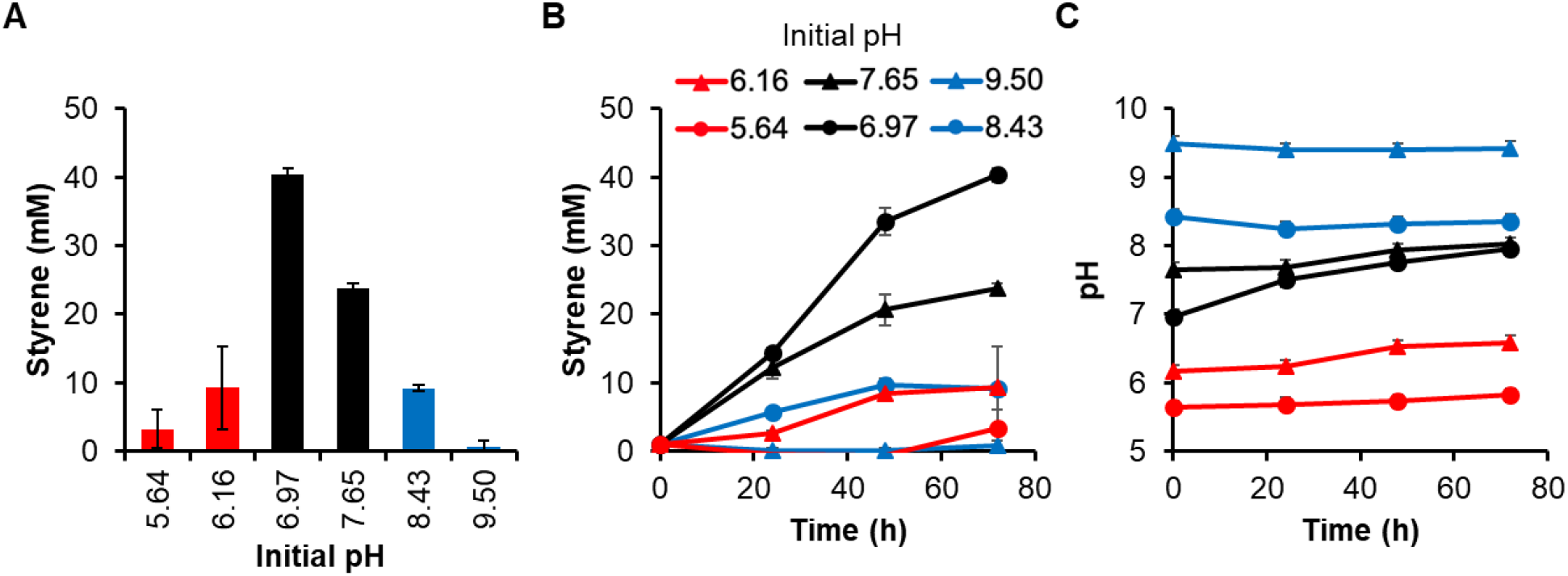
Styrene biosynthesis is highly pH dependent. Cell-free reactions containing 0.25 μM PAL2 and 1 μM FDC1 were set with several initial pH values and produced a large range of styrene titers. **(A)** Endpoint titers after 72 hours indicate a clear optimum for reactions starting at neutral pH. **(B)** The rate of styrene synthesis diminished as the pH deviated from neutral. **(C)** pH changed little over the course of the reactions. Error bars represent standard deviation of 3 technical replicates.

## 4. Conclusion

In this study, we demonstrated a substantial increase in styrene titer using a cell-free system compared to *in vivo* biosynthesis through the optimization of reaction temperature, enzyme ratios, and pH. We first determined that PAL2 and FDC1 expressed by *E. coli* CFPS were soluble and active *in vitro* and that, in combination, these enzymes produced the most styrene at 30 °C. Second, we found that cell-free styrene biosynthesis was maximized by combining a low concentration of PAL2 with a high concentration of FDC1, which would maximize conversion of *trans*-cinnamic acid to styrene. Third, we highlighted the pH sensitivity of *in vitro* styrene biosynthesis and found that reactions maintained at a neutral pH produced the highest concentration of monomer. These cell-free reactions achieved a maximum styrene titer of 40.33 ± 1.03 mM (4.20 ± 0.11 g/L), which surpasses the highest published in vivo titer of 3.36 mM (0.350 g/L) by more than an order of magnitude (Liu et al., 2018). Such high titers of the toxic styrene monomer are possible due to the lack of viability constraints in a cell-free system and the ability to finely tune the reaction environment, which makes this system a powerful example of cell-free biosynthesis as an alternative to traditional cell-based biomanufacturing methods. Although the optimal pH for styrene biosynthesis was within a range consistent with the optima for the PAL2 and FDC1 homologs used, the overall reaction rate appeared much slower than expected (Cochrane et al., 2004; Payne et al., 2015). Despite the decreased rate, the potential impact of this environmentally-friendly, cell-free styrene synthesis approach deserves further assessment through economic process models that consider substrate, catalyst, and product costs, including styrene monomer and polystyrene derivatives.

Looking forward, increased potential of the cell-free approach as a biomanufacturing platform could be achieved by investigating direct styrene polymerization. Styrene readily polymerizes in organic solvents upon heating in the presence of a radical initiator, such as benzoyl peroxide, due to the vinyl group (Abere et al., 1945). Replacing the dodecane overlay used for styrene extraction in this study with toluene could enhance the solubility of biosynthesized styrene as the solvent is heated. Despite the substantial increase in titer achieved using a cell-free system, these microliter-scale reactions cannot feasibly produce styrene in the large quantities currently derived from petroleum (James and Castor, 2011). Fortunately, larger scales are possible with cell-free protein synthesis scaling linearly up to 100 L (Zawada et al., 2011), although increasing the scale of biosynthesis reactions with a hydrocarbon overlay presents technical challenges. Furthermore, reaching the capacity of petrochemical plants would not be necessary since smaller scale biomanufacturing facilities would still benefit from an economy of scale for feedstocks while simultaneously reducing capital risk and market saturation (Claypool et al., 2014).

In sum, the laboratory example of high-titer styrene biosynthesis described here demonstrates the potential of cell-free systems for the production of toxic compounds that are currently produced by petroleum-based processes. Expanding the cell-free approach to producing more value-added chemicals, such as other plastic precursors and biofuels, and increasing the scale of these reactions could spearhead the development of economically viable alternatives to fossil fuel-derived chemicals.

## Acknowledgments

We graciously thank the Department of Energy (BER grant: DE-SC0018249), the David and Lucile Packard Foundation (2011-37152), and the Camille Dreyfus Teacher-Scholar Program for support. B.J.R. is an NDSEG Fellow (Award ND-CEN-017-095).

## Author Contributions

W.S.G., B.J.R., and A.K. performed the experiments and analyzed the data. A.S.K. and M.C.J. provided supervisory roles. All authors conceived experiments and wrote the manuscript.

## Supplemental Information

### Supplementary Methods

#### Metabolite Quantification by HPLC

Reactions were quenched by adding 10% w/v tricholoacetic acid (Sigma) in a 1:1 ratio. Precipitated proteins were pelleted by centrifugation at 21,000g for 10 minutes at 4 °C, and 20 μL of the supernatant was removed for analysis by high-performance liquid chromatography (HPLC). L-Phe was measured with an Agilent 1260 series HPLC system (Agilent, Santa Clara, CA) via a refractive index (RI) detector after passing through a reverse-phase Hypersil Gold column (4.6 mm x 150 mm; Thermo Fisher, USA). 5 μL from the prepared samples were injected at a total constant flow rate of 1 mL / min and temperature of 45 °C. The column was operated with water (solvent 1) and methanol plus 0.1% trifluoroacetic acid (solvent 2). The eluent began as a mixture of 95% solvent 1 and 5% solvent 2 before a linear gradient was applied over 8 minutes to reach a mixture of 20% solvent 1 and 80% solvent 2. This composition was held constant for 2 minutes before a second linear gradient was applied over the course of 4 minutes to return to its initial composition of 95% solvent 1 and 5% solvent 2.

**Supplementary Figure S1.**
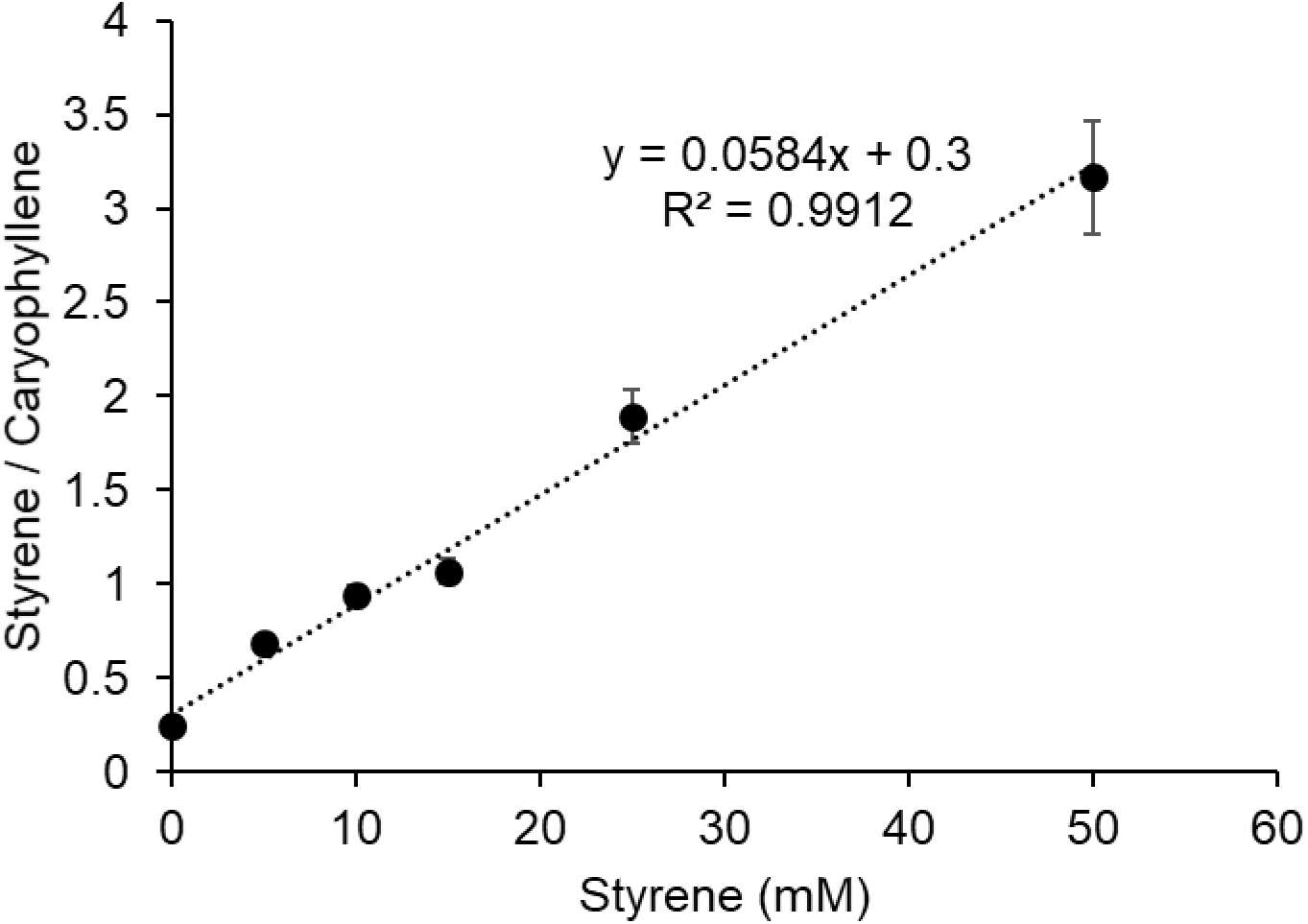
Styrene standard curve for GC-MS analysis. Dilutions of styrene dissolved in dodecane were mixed with mock cell-free reactions containing GFP in place of biosynthetic enzymes to emulate the composition of experimental samples. In standards and samples, styrene counts were normalized to the *trans*-caryophyllene internal standard in the ethyl acetate loading solvent. Error bars represent standard deviation of 3 technical replicates.

**Supplementary Figure S2.**
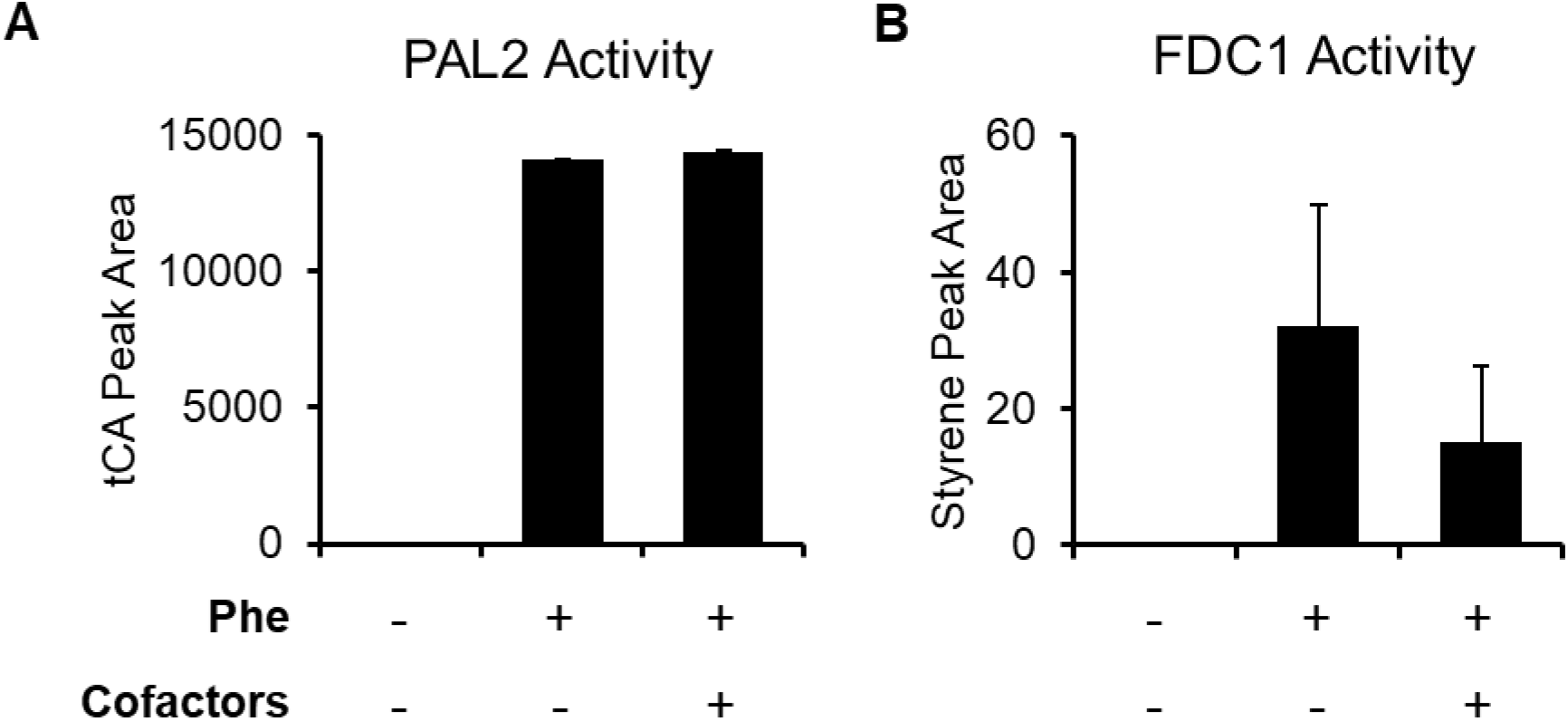
Verification of enzyme activity. **(A)** HPLC analysis indicated PAL2 activity *in vitro* is independent of cofactors, which included 1 mM ATP, NAD, and Coenzyme A. **(B)** FDC1 is active in the presence of PAL2, but styrene quantification in later experiments was performed using GC-MS for greater accuracy. Error bars represent standard deviation of 2 technical replicates.

**Supplementary Figure S3.**
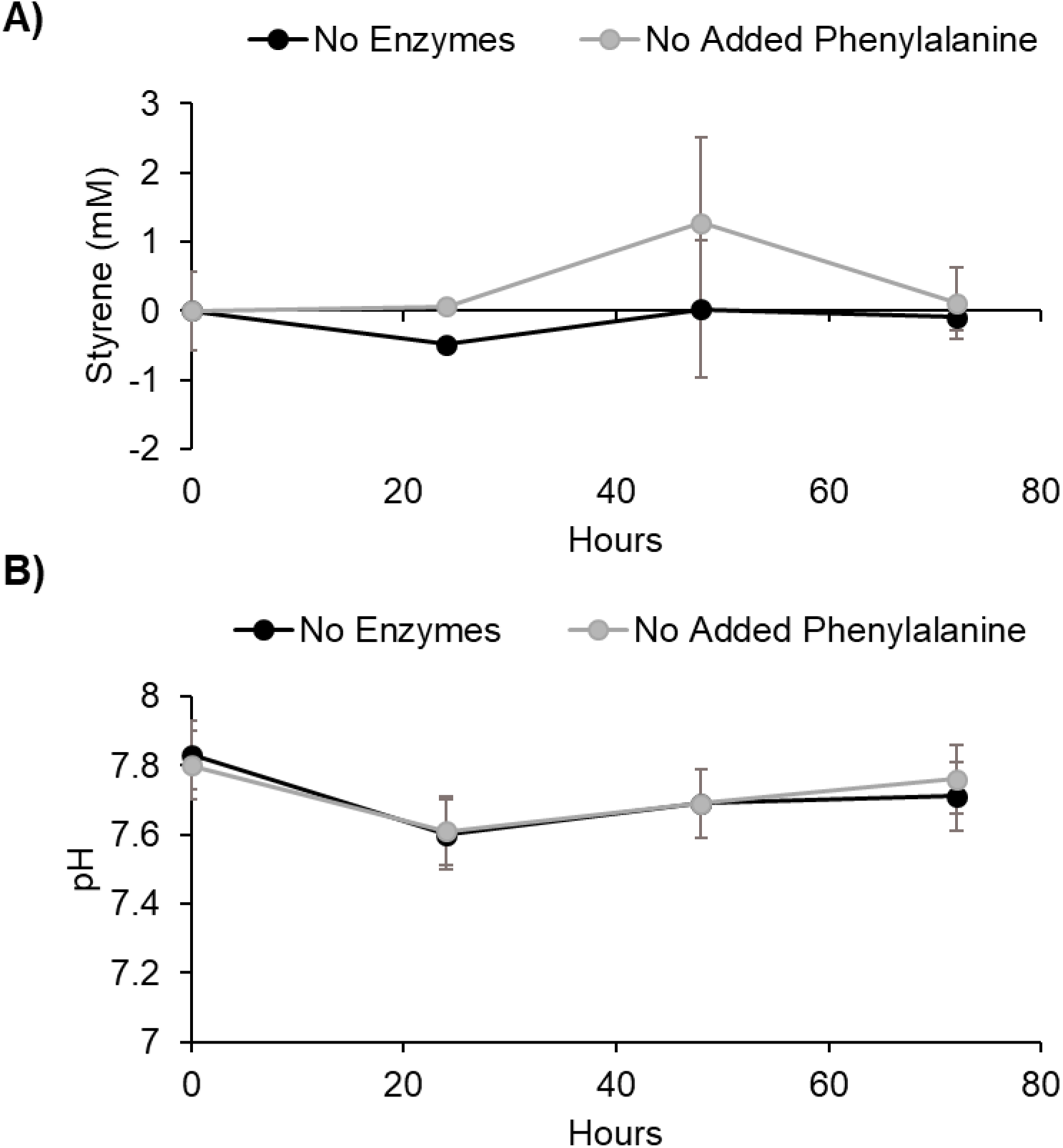
Styrene biosynthesis requires exogenous enzymes. **(A)** Reactions without PAL2 and FDC1 or additional phenylalanine do not produce styrene. **(B)** The pH of these reactions changes little over time, as expected without active glycolysis. Error bars represent standard deviation of 3 technical replicates.

**Supplementary Figure S4.**
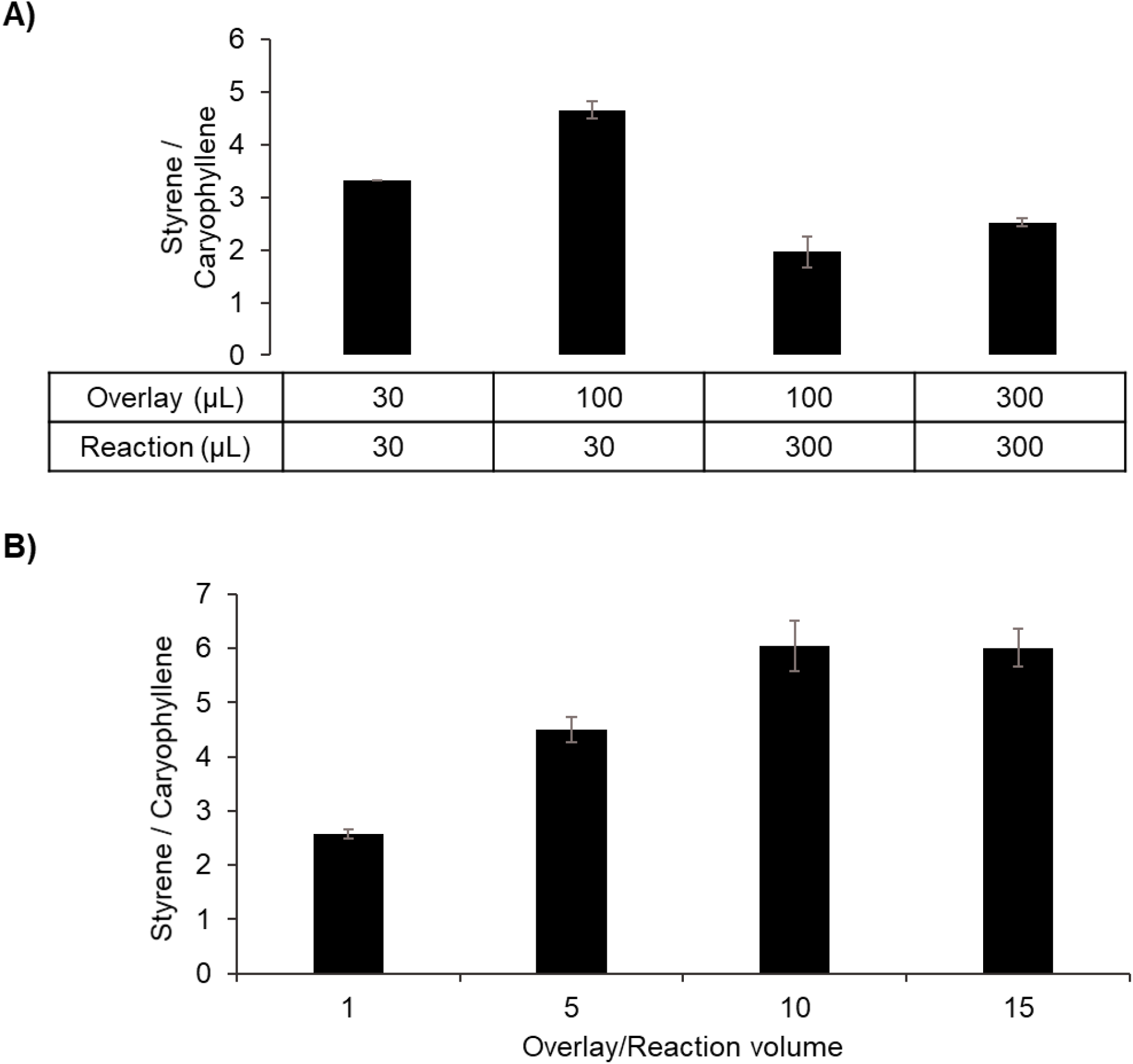
Capturing styrene with dodecane. **(A)** Due to the insolubility of styrene in aqueous solutions, a dodecane layer above the reactions was required to trap the product. Altering the reaction and overlay volumes impacted the amount of styrene captured relative to the internal standard, caryophyllene. **(B)** For 30 μL reactions, larger overlays did not inhibit styrene synthesis and captured a greater proportion of styrene per microliter sampled. These reactions contained 0.5 μM PAL2 and FDC1. Error bars represent standard deviation of 3 technical replicates.

**Supplementary Figure S5.**
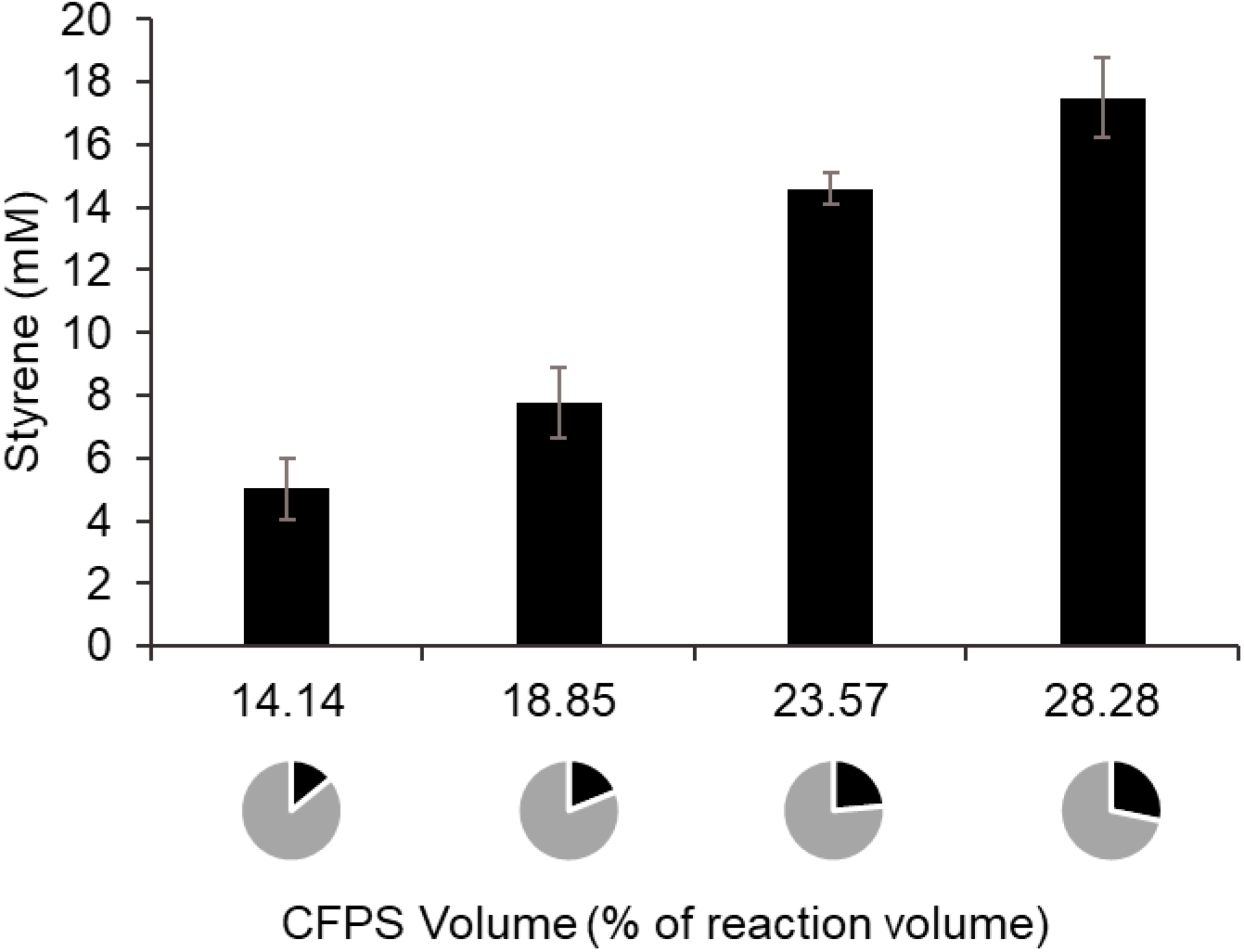
Assessing the addition of blank CFPS reactions. Reactions in Figure 4 contained various volumes of CFPS reactions without plasmid so that all reactions contained the same total volume of CFPS regardless of enzyme concentration. For reactions with 0.5 μM enzymes and 25 mM L-phenylalanine, additional CFPS volume increased final styrene titer, which accounts for the difference in titers achieved in Figure 3 (with 14% CFPS) and Figure 4 (with 28% CFPS). Error bars represent standard deviation of 3 technical replicates.

**Supplementary Table 1.**
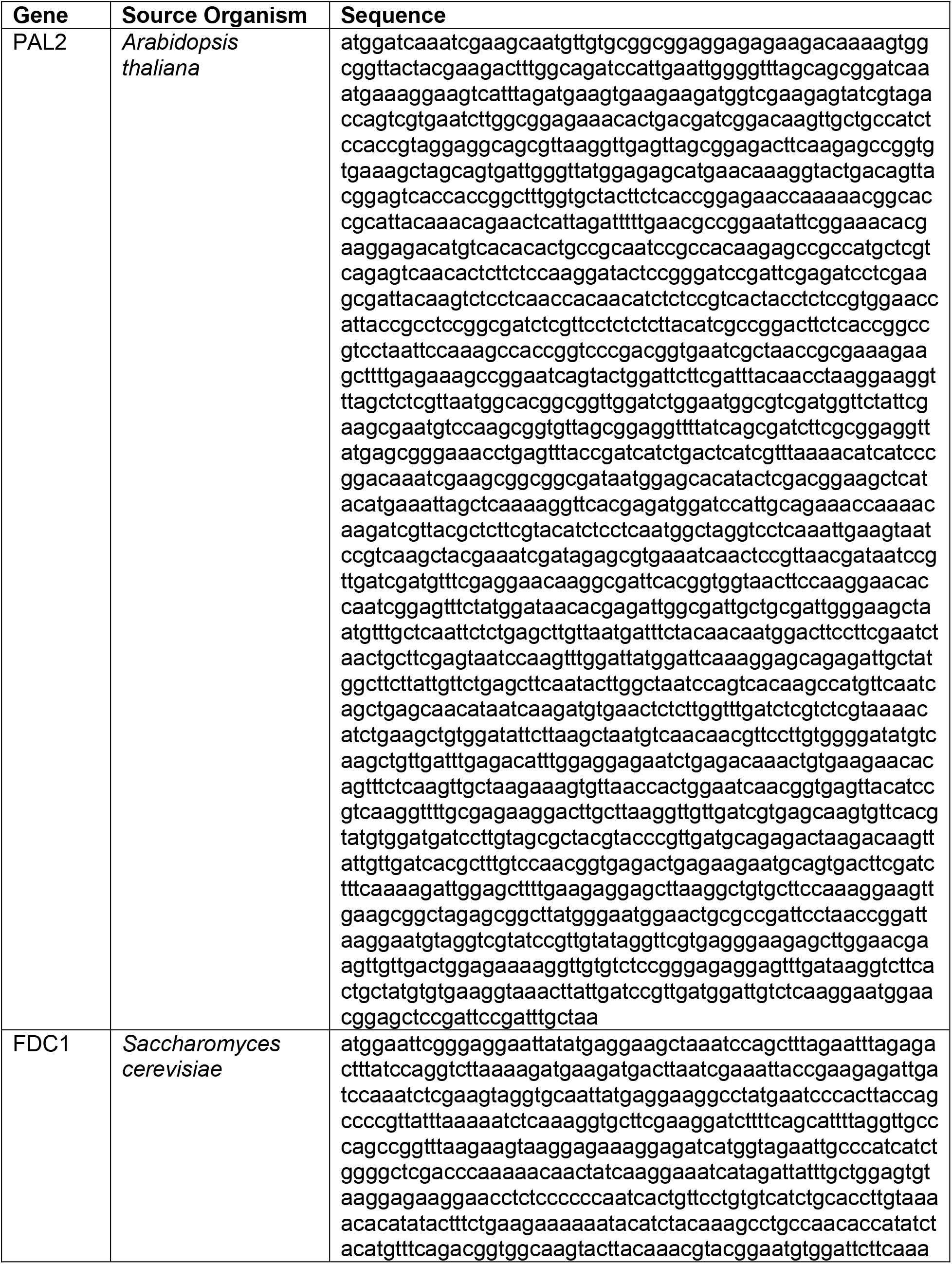
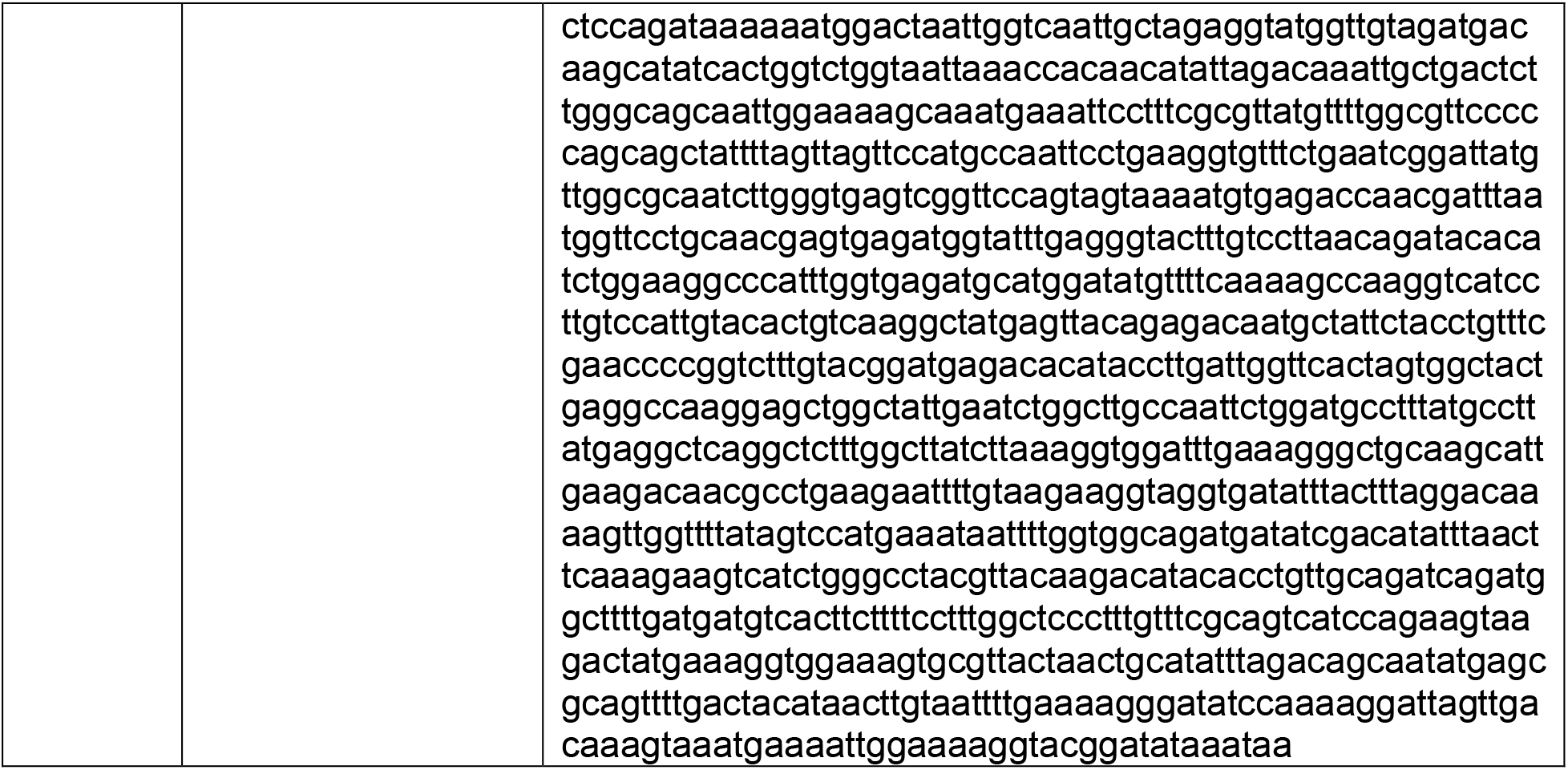
Gene sequences for biosynthetic enzymes.

